# Quantifying conformational changes in the TCR:pMHC-I binding interface

**DOI:** 10.1101/2024.08.13.607715

**Authors:** Benjamin McMaster, Christopher Thorpe, Jamie Rossjohn, Charlotte M. Deane, Hashem Koohy

## Abstract

T cells form one of the key pillars of adaptive immunity. Using their surface bound T cell antigen receptors (TCRs), these cells screen millions of antigens presented by major histocompatability complex (MHC) or MHC-like molecules. In other protein families, the dynamics of protein-protein interactions have important implications for protein function. Case studies of TCR:class I peptide-MHCs (pMHC-Is) structures have reported mixed results on whether the binding interfaces undergo conformational change during engagement and no robust statistical quantification has been done to generalise these results. It thus remains an open question if movement occurs in the binding interface that enables recognition and activation of T cells. In this work, we quantify the conformational changes in the TCR:pMHC-I binding interface by creating a dataset of 358 structures, comprising 25 TCRs, 20 MHC alleles, and 58 peptide structures in both unbound (*apo*) and bound (*holo*) conformations. In support of some case studies, we demonstrate that all complimentary determining region (CDR) loops move to a certain extent but only CDR3α and CDR3β loops modify their shape when binding pMHC-Is. We also map out the contacts between TCRs and pMHC-Is, generating a novel fingerprint of TCRs on MHC molecules and show that the CDR3α tends to bind the N-terminus of the peptide and the CDR3β tends to bind the C-terminus of the peptide. Finally, we show that the presented peptides can undergo conformational changes when engaged by TCRs, as has been reported in past literature, but novelly show these changes depend on how the peptides are anchored in the MHC binding groove. Our work has implications in understanding the behaviour of TCR:pMHC-I interactions and providing insights that can be used for modelling T cell antigen specificity, an ongoing grand challenge in immunology.

## 1 Introduction

T cells are essential cells of the adaptive immune system, responsible for identifying and eliminating foreign pathogens or malfunctioning cells to maintain homeostasis. To discriminate between foreign antigens and self-peptides, T cells use their surface bound T cell antigen receptors (TCRs) to scan linearised peptides presented by major histocompatability complex (MHC) molecules.

TCRs are hetero-dimeric molecules, consisting of an α- and β-chain (or a γ- and δ-chain for a smaller subset) with a constant region that anchors them to the cell membrane and a variable domain responsible for antigen binding [1]. The antigen binding region is formed of six protein loops (three on the α-chain and three on the β-chain) known as the complimentary determining regions (CDRs). These loops are the product of a stochastic gene rearrangement process known as V(D)J recombination that occurs during T cell development in the thymus and resulting in the huge breadth of T cell diversity [2]. Unlike their B cell counterparts, known as B cell receptors or antibodies, TCRs do not undergo any further modifications to influence their antigen specificity.

On the presentation side, class I MHC molecules are found on all nucleated cells and are responsible for displaying peptide fragments of degraded proteins on the cell surface. Based on recognition from T cells, these molecules mark the cells’ internal state as diseased or healthy, helping to ensure homeostasis through the removal of diseased cells. The class I peptide-MHC (pMHC-I) complex is formed of a membrane-anchored domain and an antigen-binding domain with a groove holding the peptide created by two α-helices and a floor formed of seven anti-parallel β-strands. The complex is stabilised by a β_2_-microglobulin protein underneath the antigen binding domain. Recognition of a pMHC-I by a T cell can start a cascade of signalling molecules leading to an immune response.

Proteins by nature can be dynamic entities; they can exist in multiple conformations and use movements to carry out specific biological functions. For example, Kinesins undergo large conformational changes when phosphorylated to “walk” down cytoskeletal structures and transport other molecules around the cell [3]. When proteins interact with other proteins, there can often be reported conformational changes to improve the selectivity and strength of binding to one another [4].

These changes in structure come with entropic and enthalpic considerations and are thought of in three modalities: (1) the “lock-and-key” model states that neither protein moves and the shapes fit together incurring little to no free energy penalty during binding, (2) the “induced-fit” model assumes that one protein moves while the other remains fixed, or (3) the “pre-existing equilibrium” (also called “conformational selection”) states that the proteins exist as conformational ensembles and that when the conformations are both right the proteins can bind [4].

Emerging reports have sought to quantify these principles in antibodies and show that they undergo some conformational change in their CDR regions when binding to antigens [5]. In past literature of TCR and pMHC-I interactions, there has been evidence of some conformational changes and plasticity [6–12]. Kjer-Nielsen *et al*. show that the CDR loops of a TCR move to form the bound complex with a pMHC-I [8, 9]. Tynan *et al*. also show that the binding of a TCR onto a pMHC-I molecule flattens a bulging peptide [12]. Contrarily, Chen *et al*. show in a different TCR:pMHC-I both the TCR and pMHC-I maintain their shape a fit together in a “lock-and-key” mode [11]. Other past studies have argued that structural rearrangements in the TCR:pMHC-I binding interface affect T cell activation and function [13]. Armstrong *et al*. analysed these early structures from the 9 unique TCR available in bound and unbound forms [14]. Their work concluded that all CDR loops undergo conformational changes but CDR3α and CDR3β have the largest movements. They also argue that TCR:pMHC interactions fall somewhere into the paradigms of “induced-fit” and “pre-existing equilibrium” protein-protein interactions. Since Armstrong *et al*.’s work in 2008, over 500 new TCR structures have been deposited in the RCSB protein data bank (PDB) [15] and little has been done to conduct a similar analysis on the larger amount of available structures. Questions remain on how past findings generalise to broader TCR and pMHC-I interactions and to quantify the degree and type of movements these molecules may undergo between the unbound (*apo*) TCR and pMHC-I complexes, and the bound (*holo*) TCR:pMHC-I complex.

In this work, we present an analysis of the conformational changes observed during the engagement of TCRs and pMHC-Is. We leverage databases containing TCR and pMHC-I structures to curate a dataset of 358 structures with 25 TCRs binding 20 MHC alleles and 58 different peptides, with both unbound (*apo*) and bound (*holo*) forms of TCRs and pMHC-Is. Using this dataset, we conduct a robust statistical analysis of the amount and type of movement these entities undergo when coming into contact with one another at a scale not previously done in the literature. Our analysis reveals that all CDR loops undergo conformational change between *apo* and *holo* states but that only the CDR3 loops are flexible. By mapping the contacts made between TCR CDR loops and the surface of the pMHC-I complex, we show that the interactions occur in a constrained space for each CDR loop and that both CDR3 loops are equally involved in peptide binding, with the CDR3α focused on the N-terminus of the peptide and the CDR3β focused on the C-terminus of the peptide. We also show that peptides can undergo conformational change when engaged by TCRs and this movement is dictated by how the peptide is anchored in the MHC binding groove. Our work provides a quantitative picture of TCR engagement with pMHC-I molecules, generating insights into the behaviour of T cell antigen recognition.

## 2 Results

We began our analysis by creating a dataset of TCRs and pMHC-Is structures in both their *apo* and *holo* forms. These structures were collected from the STCRDab [15] and histo. fyi [16] and were subject to the quality screening and alignment procedures described in Section 4.1. The TCR gene usage, MHC allele, and peptide redundancy of the dataset are visualised in Fig. 1A-C. The dataset contains 25 unique TCRs as defined by their IMGT CDRs, 20 unique MHC alleles, and 58 unique peptides.

**Figure 1:**
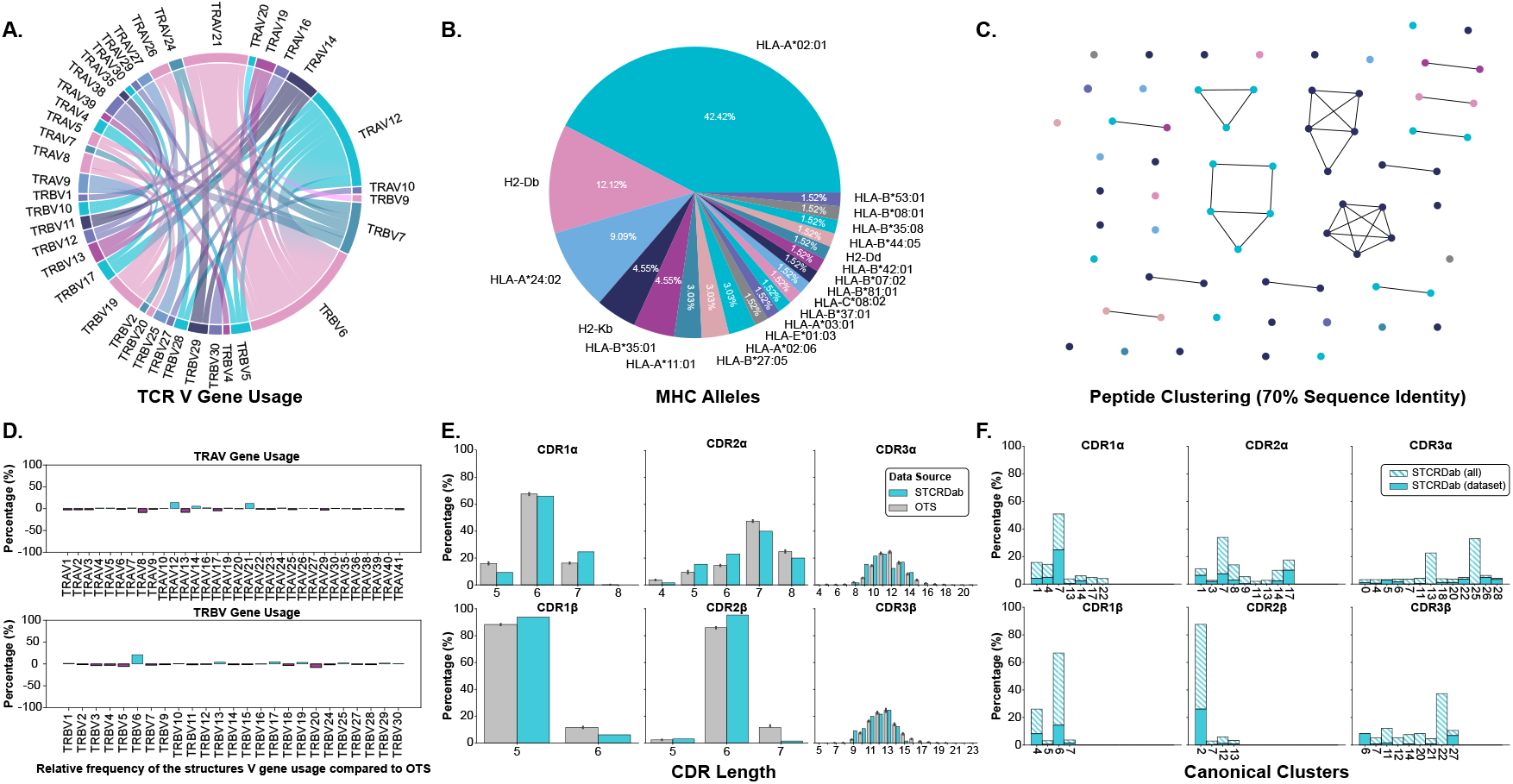
Description of TCRs and pMHC-Is in the *apo*-*holo* analysis dataset. **A**. Pairings of V genes used by the TCRs in the analysis. **B**. Proportion of different MHC alleles in the dataset. **C**. Clustering of peptides in the dataset based on sequence similarity. Clusters are formed from peptides with 70% sequence similarity and each peptide is coloured by the MHC allele presenting it according to the colouring in panel *B*. **D**. Comparison of V gene usage between the structure dataset and a background of TCRs taken from OTS. **E**. Comparison of CDR lengths between the structure dataset and a background of TCRs taken from OTS. **F**. Dataset coverage of canonical loop clusters from the whole STCRDab.

We compared the TCRs in the dataset to a background of TCRs sampled from the Observed TCR Space (OTS) database [17] to ascertain how representative the TCRs used in this analysis are to an expected distribution of TCRs. We compared common sequence properties of the molecules including V gene usage, CDR length, and amino acid composition of the CDR loops. Fig. 1D shows that for the most part, V genes are well represented within our dataset, as indicated by the near zero per cent enrichment or depletion in the plot. However, there are several notably enriched genes: TRAV12, TRAV21, and TRBV6, as well as some depleted genes: TRAV8, TRAV13, and TRBV20. Although these results indicate some bias in the dataset, some of these enrichments/depletions are expected as this work focuses solely on TCRs interacting with pMHC-I derived from CD8+ T cells. For example, TRAV12 and TRAV21 have been shown to be enriched in populations of CD8+ T cells whereas TRAV8 is comparatively enriched in CD4+ T cells, explaining its depletion here [18]. When comparing CDR lengths between the selected structures and OTS background (Fig. 1E), it seems the distributions are well matched with the top loop length matching in 5 of the 6 loop types, only CDR3α has an apparent depletion of the top OTS loop length mode. A comparison of amino acid composition in the CDR loops (shown in Fig. S1) also shows no major differences between the selected structures and the OTS background.

The representation of canonical loop classes was also assessed in the selected structures. The process for assigning loops to canonical classes is described in Section 4.3. Fig. 1F highlights the coverage of canonical classes in our dataset. 78.26% of canonical loop classes are represented in our dataset meaning most standard configurations of loop conformations are included.

Although the data contains some biases, through these results we show the dataset of structures in *apo* and *holo* conformations is representative of the broader TCR:pMHC-I interactions.

### 2.1 TCRs and MHCs undergo significant conformational changes between *apo* and *holo* states

Our analysis shows that all six CDR loops undergo conformational change when a TCR engages with a pMHC-I. Fig. 2A and B depict and quantify these changes respectively, with a dashed red line in Fig. 2B as a visual aid of the expected noise based on other reports of general noise in crystallography data [5]. Fig. 2C categorises the movements of all loops between *apo* and *holo* states. The quantification of each loop is reported as the backbone root mean squared deviation (RMSD) between *apo* and *holo* conformations (See Section 4.2). The CDR1α, CDR2α, and CDR3α loops have a mean change of 1.63 Å, 1.36 Å, and 2.21 Å respectively, with a standard deviation (SD) of 1.13 Å, 0.73 Å, and 1.67 Å. Similarly, the CDR1β, CDR2β, and CDR3β, loops have a mean change of 0.82 Å, 0.88 Å, and 1.38 Å respectively, with a of SD 0.43 Å, 0.72 Å, and 0.98 Å. Performing a Kruskal-Wallis test reveals a significant difference between the amount of conformation each loop undergoes (p-value of 3.03 × 10^−6^ at a significance level of 0.05). *Post hoc* Wilcoxon rank-sum tests with adjusted significance levels using a Bonferroni correction show that the α-chain moves more than the β-chain for the CDR1 and CDR2 loops but the same significance could not be determined for the CDR3 loops although the mean movement is higher in the CDR3α compared to CDR3β. When considering both chains together, our *post hoc* tests indicate that the CDR3 loops move more than the CDR2 loops. There was not enough statistical power to say the same thing about the CDR1 and CDR3 loops but the mean movement is higher for both CDR3 loops over the CDR1 loops.

**Figure 2:**
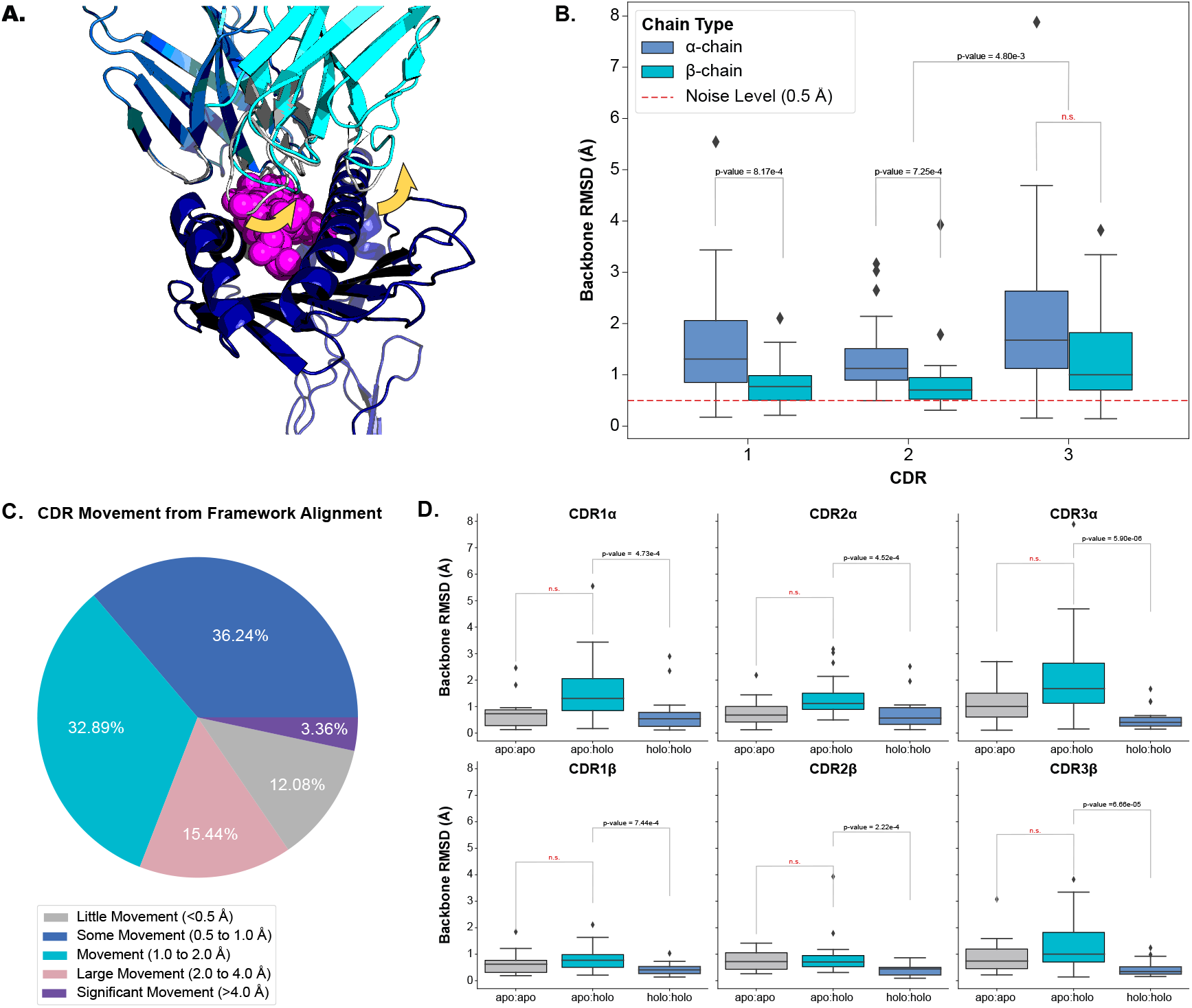
Quantifying the movement of each CDR loop. **A**. An example of CDR movement between the *apo* PDB ID 4jfh TCR (grey) and the *holo* PDB ID 4jfd TCR:pMHC-I structure (blue and cyan) as denoted by the yellow arrows. **B**. Comparison of each loop between *apo* and *holo* states. There is an overall significant difference in the amount of movement each loop undergoes based on the Kruskal-Wallis test (p-value of 3.03× 10^−6^; significance level < 0.05). **C**. Percentage of different movement categories from all CDRs. **D**. Comparison of different movement types for each CDR loop. *apo*:*apo* refers to changes between different *apo* structures of the same TCR (11 TCRs), *apo*:*holo* refers to changes between *apo* and *holo* structures (25 TCRs), and *holo*:*holo* refers to changes between different *holo* structures of the same TCR (18 TCRs). There are significant differences between the movement types based on a p-value of 4.92 *×* 10^−16^ from a Kruskal-Wallis test (significance level < 0.05).

To ascertain whether these movements are the result of an engagement between the molecules or noise between different crystal structures, we compared the changes between *apo* and *holo* structures (25 TCRs) to those between different *apo* structures (11 TCRs) and between different *holo* structures (18 TCRs) of the same TCR. Fig. 2D illustrates that there is a difference between these comparisons and performing a Kruskal-Wallis test on the comparisons yields significant results at a significance level of 0.05 (the p-value of the test is 4.92 × 10^−16^). Performing *post hoc* Wilcoxon rank-sum tests with adjusted significance levels using the Bonferroni correction shows that the changes between *apo* and *holo* structures are significantly higher than between the changes of different *holo* structures for all six loop types (our baseline used in this analysis). Although an increased trend was observed, there was not enough statistical power to distinguish the changes between *apo* structures of the same TCR and the changes between the *apo* and *holo* structures for all loops.

Our analysis shows that all six CDR loops undergo conformational change between *apo* and *holo* states and that there is some variation in the amount of change between loop types.

### 2.2 Only CDR3 loops deform when binding pMHC-Is

The results of the previous section encompass two different types of movements in the TCR: (1) bulk movements driven by changes in the anchors of CDR loops relative to the framework region and (2) loop flexibility driven by deformation in the CDR loops themselves. Again, anecdotal reports in the literature suggest differing views on the amount of flexibility CDR loops have. In some cases it has been shown that the CDR loops maintain a rigid body-like state, keeping their general shape while engaging with pMHC-I molecules [11]. Other studies suggest that all CDR loops have some type of plasticity, disrupting their canonical forms, when engaging with pMHC-I molecules [9]. In antibodies, non-CDR-H3 loops have practically no backbone deformation, and only a small amount of deformation is seen in CDR-H3 loops [5].

Here, we investigate the flexibility of TCR CDR loops to classify each loop as a rigid body, where all movement is the result of bulk movements, or as plastic entities, where flexibility also adds to the overall movement of the loop. We investigate this in three different modalities to establish robust descriptions of the deformation (Fig. 3A-D).

**Figure 3:**
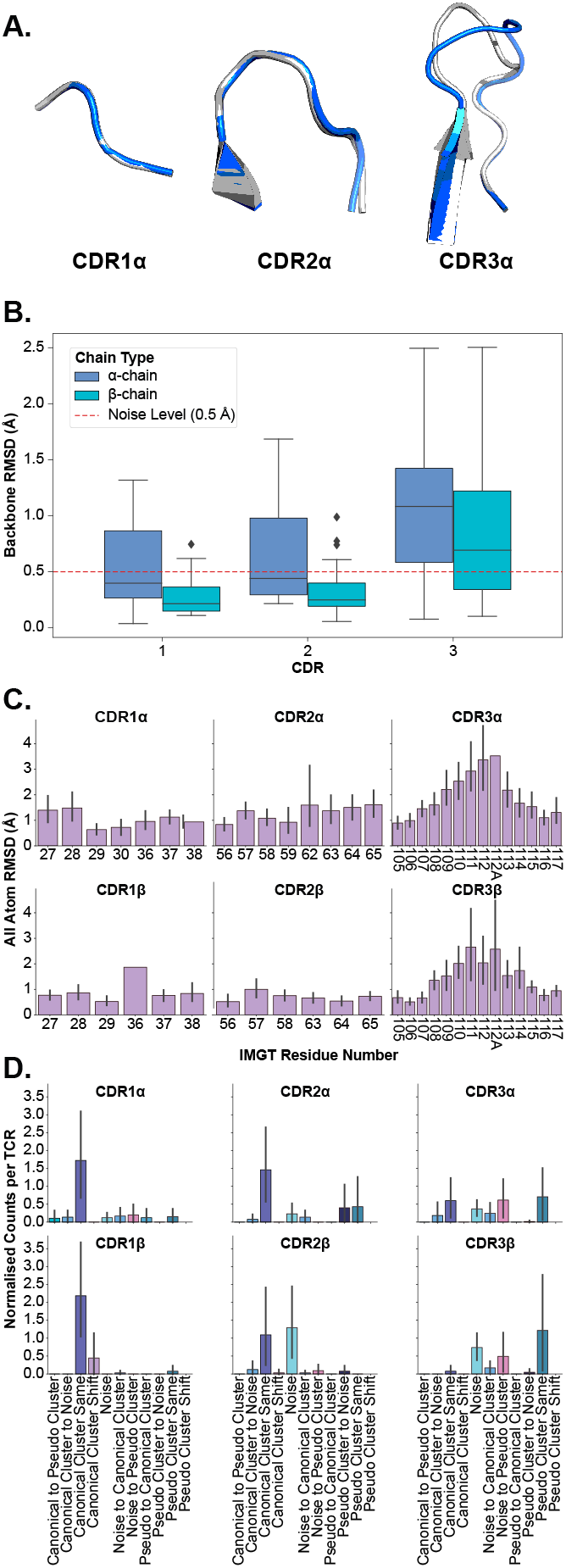
Examining the deformation of each CDR loop. **A**. An example of loop deformation in the α-chain CDR loops between the *apo* (grey; PDB ID 2bnu) and *holo* (blue; PDB ID 2bnr) states. **B**. Comparison of each loop between *apo* and *holo* states after aligning loops. **C**. Per-residue RMSD changes of each CDR loop after aligning loops. **D**. Counts of shifting cluster types between *apo* and *holo* states for each type of loop.

In the first instance, we perform the same comparison of backbone RMSD of all CDR loops (see Section 2.1) but this time aligning the loops on their backbones before measuring the differences to consider only the movement in the loops themselves. Fig. 3B depicts the quantification of these measurements. What becomes apparent is that only the CDR3α and CDR3β loops have their medians above the noise level line annotated on the figures, showing that these are the only significantly flexible loops. These findings are supported by *post hoc* Wilcoxon rank-sum tests that show only the differences between *apo*-*holo* comparisons and the *holo*-*holo* background for the CDR3α and CDR3β (see Fig. S2).

In the second case, we look at the heavy atom RMSD between *apo* and *holo* states of each residue in the loops. Fig. 3C shows that the CDR1 and CDR2 loops (from both the α- and β-chain) have uniform-like profiles where every residue moves about the same amount, whereas the CDR3α and CDR3β loops have normal-like distributions meaning the middle residues move more than those towards the anchors of the loops.

Finally, we analyse canonical clustering for each loop type and ascertain whether each kind of loop is moving from its canonical cluster group or remaining in the same structural cluster (see Section 4.3 for a description of the annotations). Fig. 3D depicts the results from this analysis. The figure shows that for CDR1 and CDR2 loops on both the α- and β-chain, the mode is to stay in the same canonical cluster between *apo* and *holo* states. For CDR3α and CDR3β, there is a larger variety of modes, including remaining un-clustered as noise, only forming canonical clusters in the *holo* state, and only clustering into pseudo clusters that remain between *apo* and *holo* states. The counts of these plots are listed in Table 1.

**Table 1:**
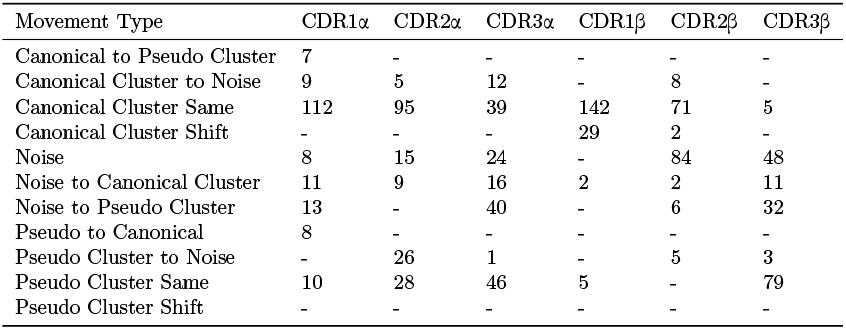
Numerical counts of the different types of cluster shifts undertaken by each loop type between *apo* and *holo* states.

| Movement Type | CDR1α | CDR2α | CDR3α | CDR1β | CDR2β | CDR3β |
| --- | --- | --- | --- | --- | --- | --- |
| Canonical to Pseudo Cluster | 7 | - | - | - | - | - |
| Canonical Cluster to Noise | 9 | 5 | 12 | - | 8 | - |
| Canonical Cluster Same | 112 | 95 | 39 | 142 | 71 | 5 |
| Canonical Cluster Shift | - | - | - | 29 | 2 | - |
| Noise | 8 | 15 | 24 | - | 84 | 48 |
| Noise to Canonical Cluster | 11 | 9 | 16 | 2 | 2 | 11 |
| Noise to Pseudo Cluster | 13 | - | 40 | - | 6 | 32 |
| Pseudo to Canonical | 8 | - | - | - | - | - |
| Pseudo Cluster to Noise | - | 26 | 1 | - | 5 | 3 |
| Pseudo Cluster Same | 10 | 28 | 46 | 5 | - | 79 |
| Pseudo Cluster Shift | - | - | - | - | - | - |

Fig. 3A illustrates these conclusions, where CDR1α and CDR2α look nearly identical between *apo* and *holo* states, but CDR3α has large changes in the middle of the loop. These results show that in general the CDR1 and CDR2 loops act as rigid bodies, deforming very little when engaging with pMHC-Is, but the CDR3 loops undergo plastic deformation to enable the interactions with the pMHC-I molecule.

### 2.3 Identifying TCR contact points on pMHC-Is

The CDR loops have been well-established as the pMHC-I binding portion of the TCR [1], but the equivalent relevant binding portion of the pMHC-I has been less studied collectively across many different structures. Thus, we set out to map out the interacting regions from the pMHC-I perspective to conduct further analyses. As described in Section 4.5, we mapped the contacts of TCR CDR loops onto the pMHC-I surface. Fig. 4A depicts these contacts on the surface of the MHC molecule and the peptide.

**Figure 4:**
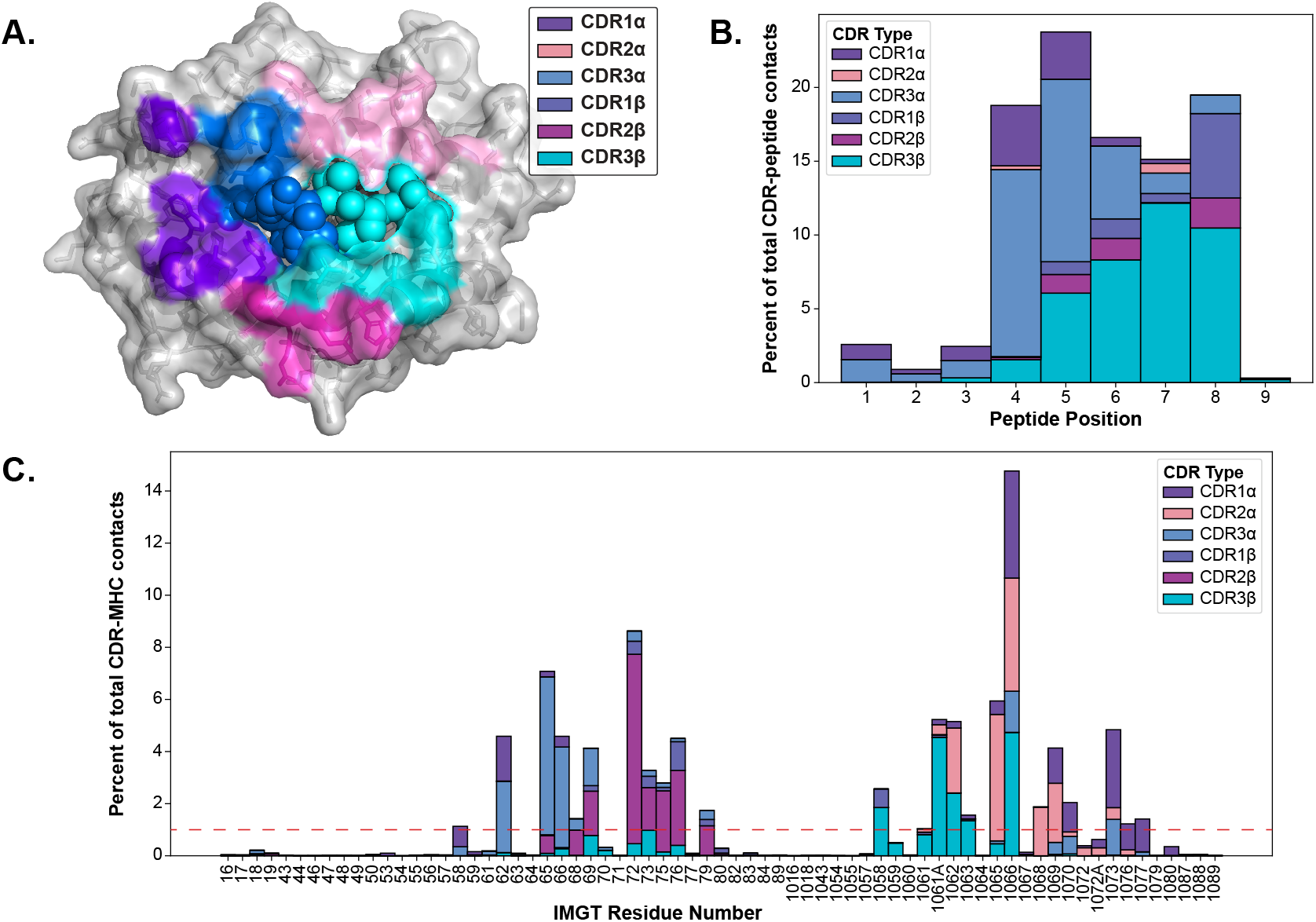
Visualizing the contacts made between TCR and pMHC-Is. **A**. The contacts made between TCR CDR loops and pMHC-Is. Here, all of the contacts (<5 Å between heavy atoms) that make up over 1% of an MHC residue contact (denoted by the red line in panel *C*) for all of the TCRs bound to pMHC-Is in the STCRDab [15] are mapped to a reference MHC molecule (PDB ID 1hhi). Notably, there are no residues dominantly contacted by CDR1β at this threshold. **B**. Distribution of contacts made between CDR loops and nonamer-peptides from the STCRDab structures. **C**. Distribution of contacts made between CDR loops and MHC molecules from the STCRDab structures.

When looking at the loops contacting the peptide (Fig. 4B), there is a trend that the CDR3α forms the majority of the contacts with the first half of peptides (p1-p5) and that the CDR3β forms majority of the contacts in the second half of the peptide (p6-p9). This description of loop engagement is in support of past literature that, despite its lack of D-gene segments, the CDR3α loop has a large structural diversity [19] and is important in determining the specificity of a TCR [20–23]. These results also show that other loops, such as CDR1α and CDR1β, can form part of the peptide binding interface as has been previously documented [24]. These trends extend to other non-nonamer peptides shown in Fig. S4.

Through this analysis, we were also able to identify the fingerprint of TCR interactions on MHC molecules and determine specific residue positions that correspond to the TCR contact points of the MHC. The conserved binding mode of TCRs on pMHC-Is has been well described in past literature [1, 24]. Fig. 4A displays the dominant loop contacting each residue and Fig. 4C shows the percentage of contacts each loop makes with the MHC residues by IMGT number.

Our generated contact maps support that both TCR chains are equally involved in contacting the peptide and each chain has a preferred side of the peptide. The contact maps also provide an alternative means to quantify the diagonal binding mode TCRs take across the MHC.

### 2.4 Peptides undergo varying amounts of conformational change based on MHC anchor locations

Establishing that TCR CDR loops undergo conformational changes when binding pMHC-Is, we set out to investigate the changes pMHC-Is undergo between *apo* and *holo* states on the other side of the complex. Fig. 5A shows that the majority of the conformational changes happen in the peptide, and that the MHC molecule undergoes little structural movements between *apo* and *holo* forms. When comparing the peptide, MHC TCR contact positions (as described in Section 2.3), and non-TCR contact positions on the antigen binding domains of the MHC I molecules, there is significantly increased movement of the peptide. Performing a Kruskal-Wallis test shows significant differences (at a 0.05 significance level) between the regions, and *post hoc* Wilcoxon rank-sum tests with Bonferroni corrections show the peptide has an increased movement over all MHC domains. There is no significant difference between the TCR contact positions and other positions in the antigen binding domains not involved directly in TCR interaction.

**Figure 5:**
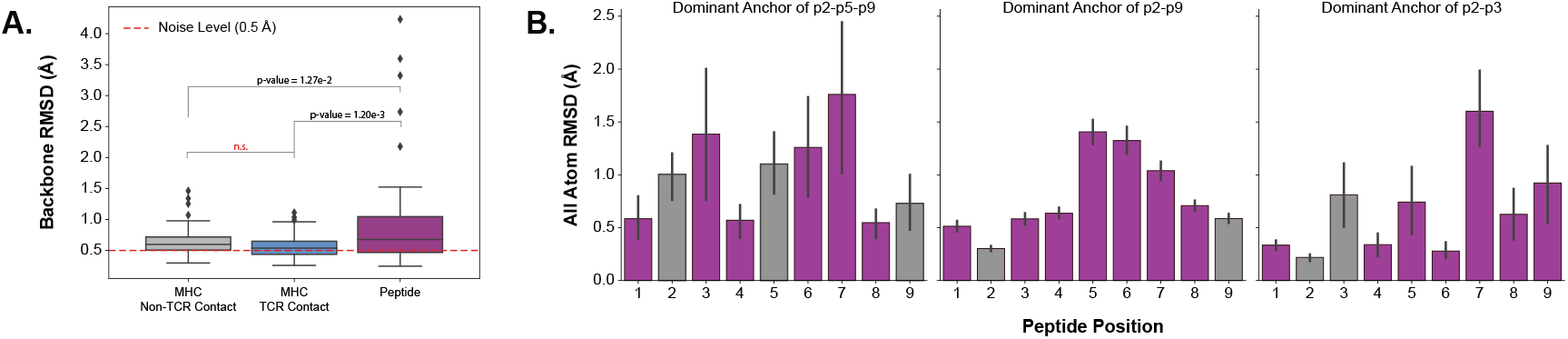
Quantifying the movement of pMHC-Is between *apo* and *holo* states. **A**. Comparison of different parts of the pMHC-I complex between *apo* and *holo* states. There is a significant difference between these components as reported by a p-value of 3.18 × 10^−4^ from a one-way ANOVA test. **B**. Effects of different peptide anchoring strategies on the deformation pattern of each peptide residue between *apo* and *holo* states.

Investigating the movement in peptides further, we found that the anchoring strategy employed by the MHC molecule dictates how a peptide will move when engaged by a TCR. We describe our procedure for identifying MHC anchors in Section 4.6. Fig. 5B shows the three different types of anchoring modes, and the results on the peptide conformational changes between *apo* and *holo* states. These plots show, that when a peptide is anchored in positions p2 and p9, as is common with the allele-specific motifs that comprise the majority of the structural data, the distribution of movement is unimodal-like where the farthest points from anchors in the middle of the peptide have the largest RMSD changes. When peptides have an additional anchor in the p5 position, the distribution becomes bi-modal-like, meaning the movement is restricted in the middle of the peptide and compensated for by freedom between anchoring residues. When the peptides are strongly anchored in positions p2 and p3 at one end of the peptide, we see a uni-modal-like distribution skewed to the peptides’ C-terminus.

These results show that the peptide is the component of the pMHC-I interface that undergoes the most change between *apo* and *holo* states and that these changes are dependent on how the peptide is anchored in the MHC binding groove.

## 3 Discussion

In this analysis, we have demonstrated the conformational changes occurring between interacting αβTCRs and pMHC-Is molecules. In particular, we showed that all six CDR loops undergo statistically significant movement between *apo* and *holo* states, but only the CDR3α and CDR3β loops have plasticity when engaging with these molecules. We also map the interactions between TCRs and pMHC-Is, highlighting the importance of CDR3α and CDR3β in making contacts with the peptides. Finally, we show that the peptides also undergo conformational changes but these changes are restricted by the way the peptides are anchored in the MHC binding groove. Our results generalise the phenomena anecdotally reported in past literature to all TCR:pMHC-I interactions, yield new insights into the TCR:pMHC-I binding, and provide considerations for modelling of TCR:pMHC-I structures and T cell antigen specificity.

With recent advances in machine learning applied to protein structure prediction [25], many new approaches have been developed to predict the structures of adaptive immune receptors from sequence to allow repertoire scale analysis of immune receptor structures [26]. Our results show important considerations for these structural prediction methods that currently do little to consider differences in the conformation of immune receptor CDRs. Here, we show that not only is there a significant difference between the *apo* and *holo* states of TCRs but also that there is movement between different *apo* states of the same TCR (see Fig. 2D). These differences may contribute to the “poor” performance of structure prediction methods at predicting CDR loops [27]. Further, in Fig. 2B, we show that the α-chain has a significantly larger movement than the β-chain CDR1 and CDR2 loops and although there is not enough statistical power to support this, the CDR3α loop has a larger change than the CDR3β loops. This fits with the narrative of recent reports of larger structural diversity in the CDR3α than the CDR3β when predicting repertoires of TCR structures [19] and results of benchmarking studies that show the CDR3α loops are harder to predict than the CDR3β loops [27].

Other works have taken similar approaches to ours in mapping out either TCR contacts with peptides [28] or TCR contacts with class I MHCs [29]. These contact maps provide useful CDR loop-specific pseudo-sequences of the MHC surface that contact TCRs. Training predictive models on these structurally constrained sequences may improve the ongoing challenge of predicting T cell antigen specificity [30]. Other works have illustrated the value of this type of information to model the MHC restriction of TCR repertoires [31].

We compared our conclusions with recent literature conducting a similar study in antibodies [5]. Antibodies have similar variable region structures to TCRs with two chains (heavy, H, and light, L) and six CDR loops forming the antigen binding domain of the molecule. How-ever, these immune cell receptors target protein antigens directly, without the need for peptide linearisation and presentation. Liu *et al*. report that the conformational changes are confined to the CDR-H3 loops, both in large-scale movements and plasticity, the other five loops undergo very little conformational change. Liu *et al*.’s results corroborate past literature on how antibody loops behave, where the CDR-H3 loop drives specificity and the other CDR loops act as stabilisation for the H3 loop [32]. In contrast, our work shows that both CDR3α and CDR3β have peptide-directed movement and plasticity, and the other CDRs also have movement that may or may not be involved in the stabilisation of CDR3 loops. This difference between receptor types may result from the fundamental differences between TCR and antibody binding of antigens [22]. However, we draw similar conclusions to Liu *et al*. that the CDR1 and CDR2 loops are mostly found in the same canonical cluster group between *apo* and *holo* states.

Through this work, we have focused on understanding the dynamics of TCR:pMHC-I interactions, but counterintuitively, we have used static structures that offer only a “snapshot” of TCR state to infer these dynamics. The reason for this is the availability of data from static X-ray crystallography is much higher than any alternatives for TCR:pMHC-I interactions. There is only one solution nuclear magnetic resonance (NMR) structure and only eight structures resolved from cryogenic electron microscopy (cryo-EM) in the STCRDab [15] with little comparison between *apo* and *holo* states. These alternative structural biology techniques may provide the field with more ways to study the dynamics of TCR:pMHC-I interactions as they are conducted in more native-like solution environments. The effects of crystal packing on the *apo* conformations in X-ray crystallography data have been discussed in other works and possibly contribute to differences in conformational states between *apo* structures [33]. Using solution NMR, Hawse *et al*. showed the dynamics of the CDR3β loop of a TCR after binding to pMHC-I and highlight the overall reduction of motility of the other CDR binding loops. Other experimental techniques such as hydrogen-deuterium exchange have been used in past works to measure pMHC-I flexibility [35–37] as well as the rigidification of TCRs when binding pMHC-I [38]. We thus hope that our investigation of TCR:pMHC-I conformational changes further prompts the study and data collection of the dynamics of these interactions.

Outside of experimental approaches, molecular dynamics (MD) simulations offer a computational alternative for exploring the dynamics of protein interactions. Several groups have sought to understand similar questions regarding the dynamics of TCR:pMHC-I interactions using these simulations. Tripathi & Salunke explored the conformational changes of the IG4 TCR in complex with the tumour epitope NYESO peptide (SLLMWITQC) [39]. Their work closely supports the results here, showing that most of the conformational change occurs in the CDR3 loops and that the CDR loops of the βchain move less than those of the α-chain. They state that these findings support the paradigm of induced fit occurring between TCRs and pMHC-Is. Other groups have focused on the conformational changes of pMHC-Is when engaged by TCRs. MD simulations show that the peptide and MHC molecule greatly affect each other’s flexibility. Hawse *et al*. show that the peptide amino acid composition modulates the MHC molecules flexibility [35] and Pöhlmann *et al*. show that MHC polymorphisms affect the flexibility of peptides, both having implications in TCR specificity [41]. Many works show that both the TCR and pMHC-I undergo a rigidification after binding to one another [42–44] and correspondingly that the CDR loops, peptide, and MHC α2-helix are more motile in the *apo* state providing evidence for pre-existing equilibrium binding between these molecules [45]. These results would be difficult to validate with X-ray crystallography data alone as it is challenging to capture the full range of protein motion from these snapshots. Finally, Alba *et al*. illustrate the uniqueness of TCR:pMHC-I interactions through MD simulations, showing the hydrogen bonding and conformational effects of a peptide are unique to the interacting TCR [46]. These help validate our lack of statistical significance between comparisons of *apo* and *holo* pMHC-Is and comparisons of *holo* pMHC-I forms with different TCRs (see Fig. S5) as, based on Alba *et al*., there is an expected heterogeneity in *holo* pMHC-I conformations. Overall, past work using MD simulations closely supports our results and provides more insight into the dynamics of TCR:pMHC-I interactions.

The introduction outlined three paradigms for protein-protein interactions: “lock-and-key”, “induced-fit”, and “pre-existing equilibrium”. Here, we have provided evidence that conformational change is a key characteristic of TCR:pMHC-I molecules, meaning both “induced-fit” and “pre-existing equilibrium” seem plausible to describe the interactions. To discriminate between these modes, further work would need to be done. “Preexisting equilibrium” could be determined by studying the unbound forms of these molecules and seeing if the *holo* states of these molecules are found within the range of *apo* forms. With limited data on this, further data collection using techniques such as NMR and cryo-EM of *apo* TCRs or pMHC-Is would be essential. In both “induced-fit” and “pre-existing equilibrium” paradigms, the interactions incur an energetic penalty as some structural rearrangements would be required for an interaction to occur. However, the flexibility of proteins can be seen as an evolutionary advantage as it allows for broader specificity of interactions from fewer stochastic sequence rearrangements events [4, 47]. Thus, the conformational changes observed here support the growing evidence that not only does our immune system rely on sequence diversity for protection from harm-causing pathogens, but structural diversity also plays a role [19].

A limitation throughout this study is the availability of non-redundant crystal structures and representation to the αβTCR and pMHC-I repertoire as a whole. Although there were several hundred structures in the dataset, careful normalisation was required to not bias the results towards overly representative TCRs or pMHC-Is. Performing more crystallography work to increase the size and diversity of the available datasets would be an obvious solution, but a highly resource-intensive endeavour. In earlier work, we have discussed the ability of machine learning models for structure predictions in overcoming some of the challenges in producing large amounts of structural data for a broader analysis of TCR and pMHC-I repertoires [26]. We speculate that this may be a promising way to overcome the biases and limitations of currently available crystallography data in the TCR:pMHC-I field, however, others have reported that these methods have limitations in predicting novel CDR shapes [48].

## 4 Methods

### 4.1 Curating *apo* and *holo* TCR and pMHC-I structures

The *apo* TCR structures and the *holo* TCR:pMHC-Is were collected from the STCRDab [15] and the *apo* pMHC-Is structures were collected from histo.fyi [16] on April 17th, 2024. The structures were all unbound αβTCRs, unbound pMHC-Is, or αβTCRs bound to pMHC-Is. Structures with a resolution greater than 3.50 Å or missing residues in the TCR variable domain or the peptide were removed from the dataset. The unbound TCRs and pMHC-Is were matched to the TCR:pMHC-I complexes based on the TCRs CDR sequences or the pMHC-Is peptide sequence and allele name. Only data points with both an *apo* and *holo* form were kept in the dataset. The resulting dataset contains 358 structures coming from 255 PDB entries and encompassing 25 unique TCRs, 20 MHC alleles, and 58 peptides. The exact structures are listed in Table S1.

All of the selected structures were renumbered using the same version of ANARCI [49] to provide consistent IMGT numbering. For every *holo* TCR:pMHC-I complex, the *apo* TCRs were aligned to the *holo* TCRs on their framework regions and the *apo* pMHC-Is were aligned to the *holo* pMHC-Is on the strands forming the floor of the MHC binding groove. For some of the calculations, *holo* structures were aligned to each other in the same manner where one of the *holo* forms is aligned on either the TCR framework region or the floor of the MHC binding groove. This created the final datasets used for calculations in the rest of the analysis and these aligned structures are provided as part of the provided code (see *Code Availability*).

### 4.2 Measuring conformational changes between states

The difference in states was measured using RMSD throughout the analysis. When larger entities were being compared, for example, the entire CDR loops, the peptide, or parts of the MHC antigen binding domain, the measure was taken using the backbone atoms (N, C_*α*_, C, O) of these entities. When comparing residues to one another, the RMSD was taken for all heavy atoms (non-hydrogen) to include information about the amino-acid side chains. Alternative measures including measuring the difference of the centre of mass of heavy atoms in each entity, the distance between C_*α*_ positions, and the difference in χ-angle were used in other plots seen in the Supplementary Information (see Fig. S3). For all analyses, the results were normalised by the type of TCR or pMHC-I being compared. This was done by taking the mean of all results for that entity type before plotting or computing statistics so that over-represented TCR or pMHC-I structures did not bias the results.

### 4.3 Clustering CDR loops based on structure

Clustering of loop structures and annotation of canonical forms was done similar to previous works by Wong *et al*. and Greenshields-Watson *et al*. [48, 50]. All CDR loops of the αβTCRs with a resolution of 3.50 Å and below were taken from the STCRDab [15]. For every loop of the same type (CDR1α, CDR2α, CDR3α, CDR1β, CDR2β, CDR3β), a pairwise distance matrix was created following the same procedure: the two loops being compared were aligned on the backbone of the five anchor residues flanking each side (N and C terminus) of the loops. The distance between their backbones was computed using the length-independent distance measure of dynamic time warping (DTW) [51]. The HDBScan clustering algorithm [52] with a minimum cluster size cutoff of five was used on the distance matrices to create clusters of similar loop structures. Clusters were assigned as canonical clusters if they contained more than two unique sequences, otherwise, they were labelled as pseudo clusters following the approach of Wong *et al*. [50].

### 4.4 Sampling TCRs from OTS

We sampled OTS [17] to generate a background of TCRs sequences to which we could compare the structures used in this analysis. All of OTS was downloaded on July 23rd, 2024. We selected all six CDR sequences and the TRAV, TRBV, TRAJ, and TRBJ gene identities based on the assigned call and removed redundant entries. The resulting dataset was then uniformly sampled 10 times, each containing a 1000 unique TCRs. This created a dataset of representative TCRs that we could compare against our selected structures.

### 4.5 Mapping contacts between TCRs and pMHC-Is

The contacts made between TCRs and pMHC-Is were determined by finding all heavy atoms (non-hydrogen) that were less than 5 Å apart between the two structures. All αβTCRs contacting pMHC-Is presenting peptide antigens with a resolution under (and including) 3.50 Å in the STCRDab (as of April 2024) [15] were considered.

### 4.6 Incorporating MHC Motif Atlas data to identify MHC peptide anchors

To determine the anchoring strategy of each pMHC-I, we annotated our datasets with information from the MHCMotifAtlas [53]. For each MHC allele in the MHCMotifAtlas, we created a simplified peptide motif based on the proportion of amino acid usage at each peptide position. These amino acids were classified as “dominant” residues if they were at over 60% of the observed amino acids in that position, “high” if they were between 30% and 60% of the observed amino acids, “medium” if they were between 20% and 30% of the observed amino acids, “low” if they were between 10% and 20% of the observed amino acids, and “very-low” if they were below 10% of the observed amino acids. Where there were “dominant” or “high” amino acids, it was assumed that these residues corresponded to residues necessary to anchor the peptide in the binding groove. Using these anchor residues, we annotated our dataset with anchors at the positions where peptide residues matched the anchor residues, the peptide lengths matched the observed simplified motif lengths, and the MHC alleles were the same. These annotations created three distinct groups corresponding to MHC alleles that anchor the peptide in positions 2 and 9 (p2-p9), anchor at positions 2, 5, and 9 (p2-p5-p9), and anchor at positions 2 and 3 (p2-p3). Where only anchors at position 2 or position 9 were found (a minor subset of the data), we added these to the p2-p9 group as it was assumed that the peptide would be anchored by another amino acid type at the missing end.

## Code Availability

All of the code, data, and notebooks used to conduct this analysis can be found at https://github.com/benjiemc/tcr-pmhc-interace-analysis.

## Acknowledgements

This work was supported by funding from the UK Medical Research Council grant number MC_UU_12010/3 to H.K., the UK Medical Research Council grant number MC_UU_00008 to B.Mc., an ARISE Fellowship from the European Union’s Horizon 2020 Research and Innovation Programme under the Marie Skłodowska-Curie grant agreement number 945405 to C.T., and the Natural Sciences and Engineering Research Council of Canada.

We would like to thank A. Greenshields-Watson, M. Raybould, F. Spoendlin, and N. Quast for their insightful conversations and expertise in some technical implementations throughout the project. We would also like to thank the developers of PyMol, Pandas, NumPy, Matplotlib and Seaborn for their contributions to the open-source community. These tools and packages were used to conduct the analysis and generate the figures for this work.

## Author Contributions

B.Mc. conceived the content, conducted the study, and wrote the article. C.T. contributed to the project conception and analysis, provided data for the analysis, and took part in the supervision of the work. J.R. contributed to certain parts of the analysis and reviewed and edited the article. C.M.D. supervised the research and reviewed and edited the article. H.K. conceived the content, supervised the research, and reviewed and edited the article.

