## Supplementary Information for "Quantifying conformational changes in the TCR:pMHC-I binding interface"

### S1 Correlation between conformational change and affinity

We were interested in whether there was a correlation between the amount of conformational change in the binding interface of TCR:pMHC-Is between *apo* and *holo* states and the affinity of the interaction. We combined the results of our analysis of crystal structures with affinity measures available in the ATLAS dataset [3]. Shown in Fig. S6 and Fig. S7, we plotted the changes in conformation versus the affinity values and measured the correlation. Only the CDR2α, CDR3α, and CDR3β loops showed a small correlation to affinity ($R^2 \geq 0.05$). The other loops show very little to no correlation. On the pMHC-I side, neither the movement in peptide, major histocompatability complex (MHC) TCR contacting regions, nor the rest of the MHC antigen binding domain showed any correlation to affinity ($R^2 < 0.05$). This analysis suggests that although there are slight correlations

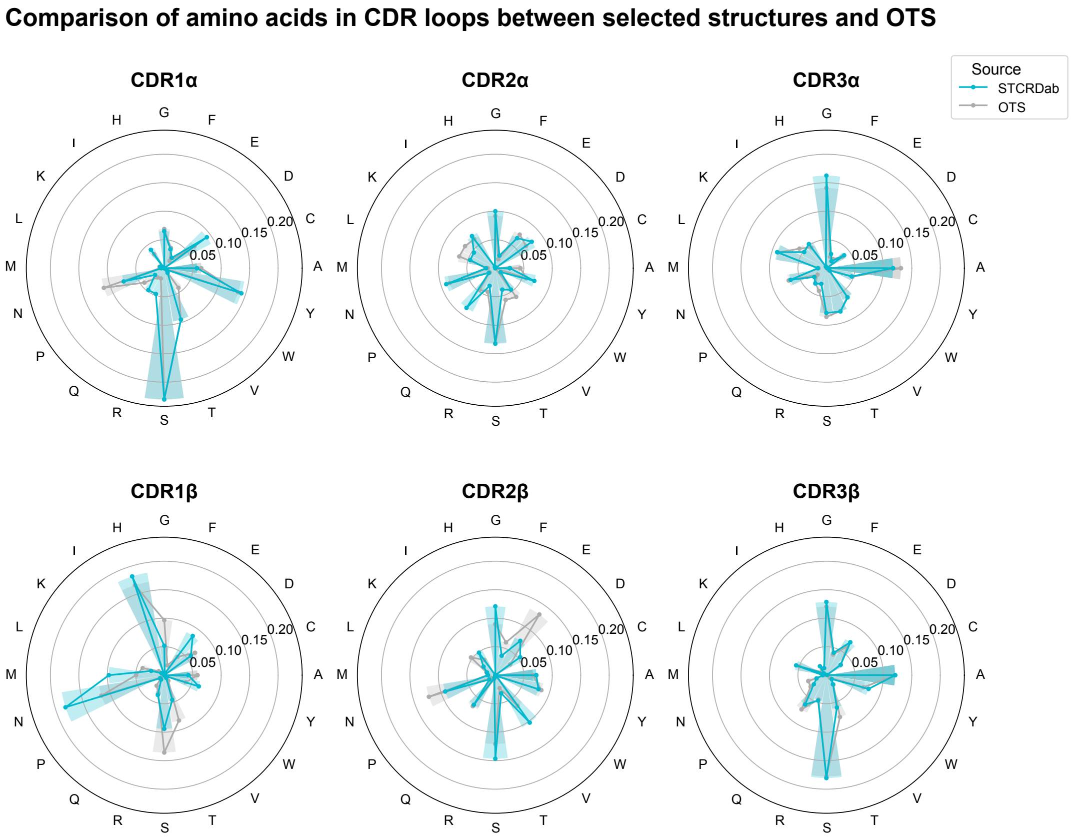


Figure S1: Comparison of the composition of amino acid residues in each CDR loop between the structures in this analysis and a comparative background of TCRs randomly sampled from OTS [1].

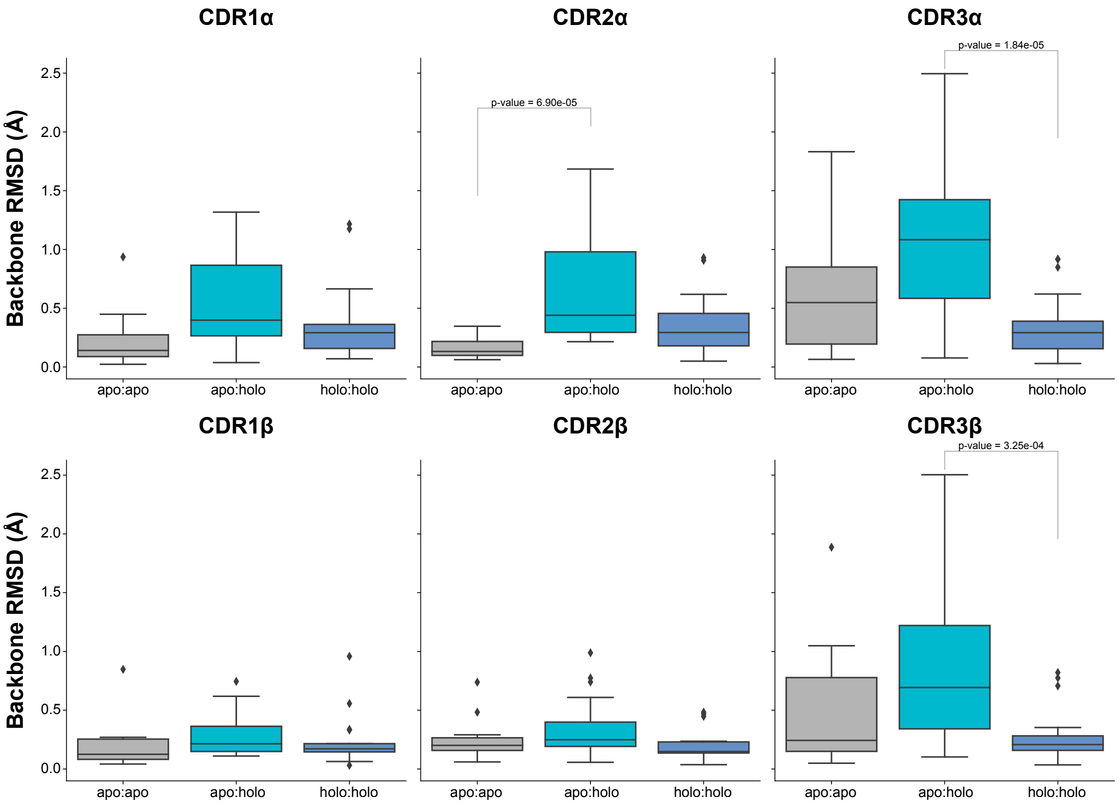


Figure S2: Comparison of different movement types for each CDR loop after alignment. *apo*:*apo* refers to changes between different *apo* structures of the same TCR, *apo*:*holo* refers to changes between *apo* and *holo* structures, and *holo*:*holo* refers to changes between different *holo* structures of the same TCR. There are significant differences between the movement types based on a p-value of $4.93 \times 10^{-15}$ from a Kruskal-Wallis test (significance level $< 0.05$). Significant *post hoc* results have been added to the plot, but non-significant test results are not shown.

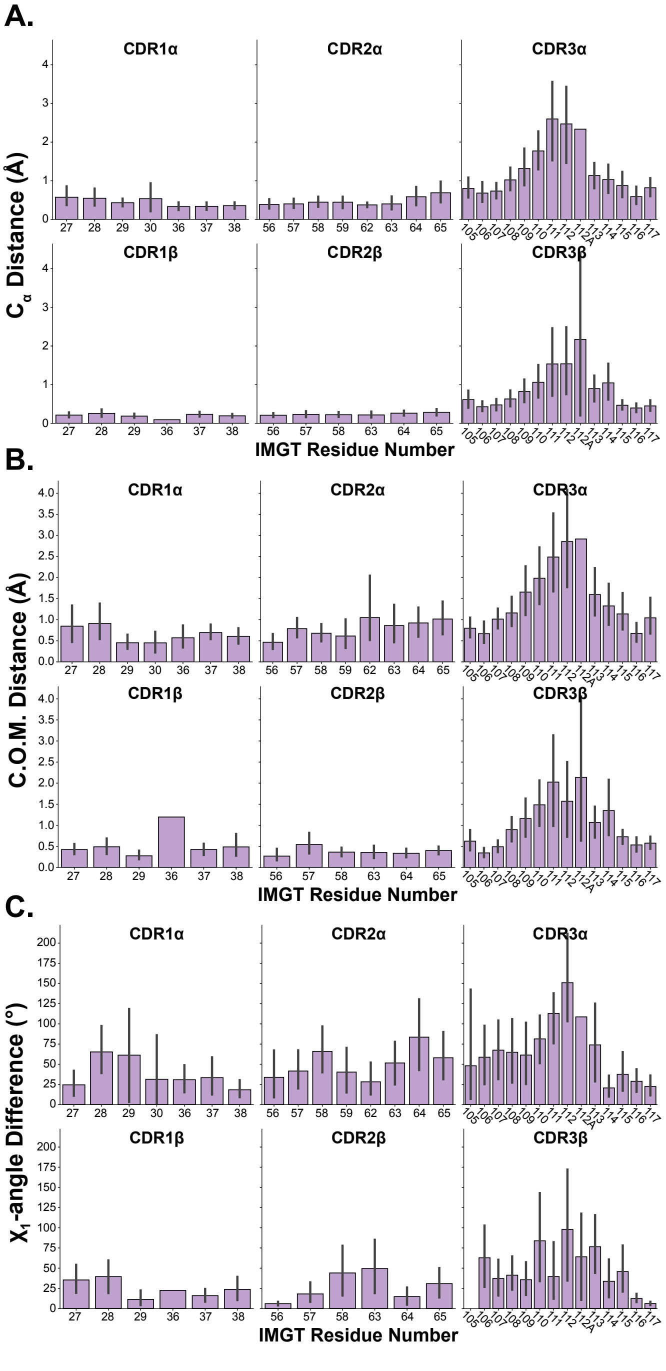


Figure S3: Describing the movement of each CDR loop in more detail. **A.** $C_\alpha$ movement between *apo* and *holo* conformations. **B.** Residue centre-of-mass changes between *apo* and *holo* conformations. **C.** $\chi_1$-angle changes between *apo* and *holo* conformations.

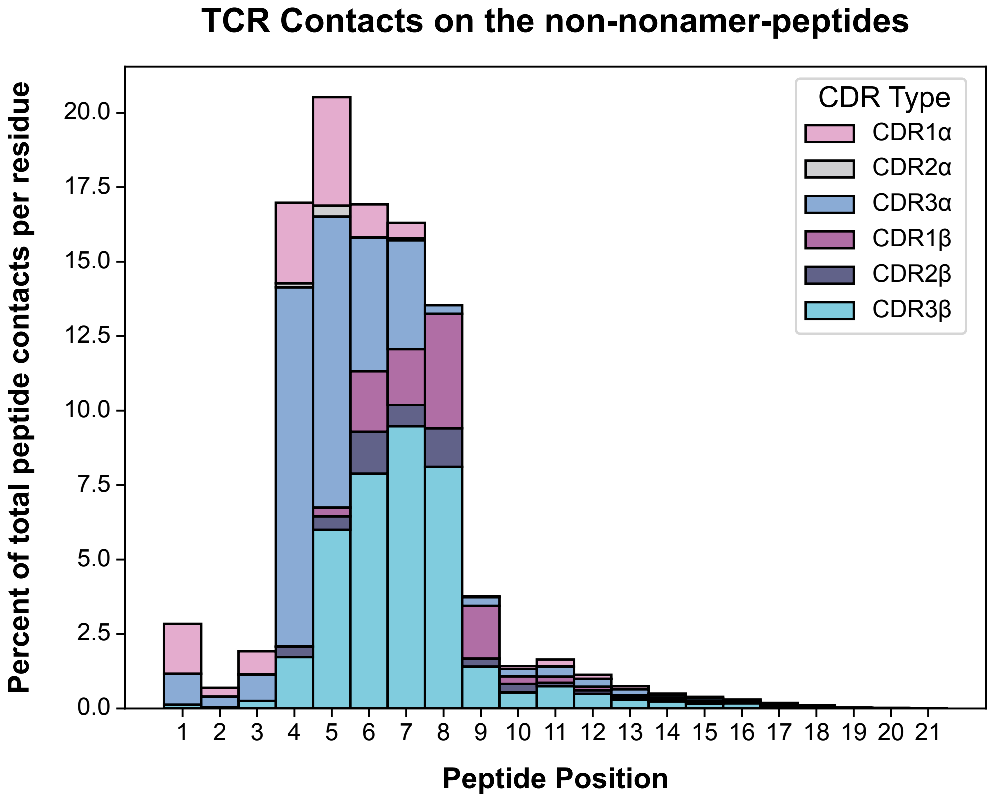


Figure S4: Distribution of contacts made between CDR loops and non-nonamer peptides from the STCRDab [2] structures.

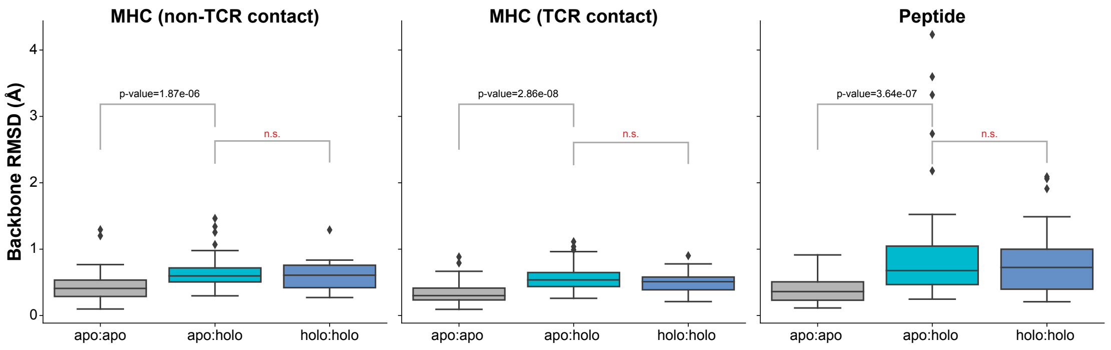


Figure S5: Comparison of *apo*:*apo*, *apo*:*holo*, and *holo*:*holo* changes for pMHC-Is. Significant differences exist between the different comparisons based on the results of a Kruskal-Wallis test at a 0.05 significance level (p-value $6.39 \times 10^{-16}$).

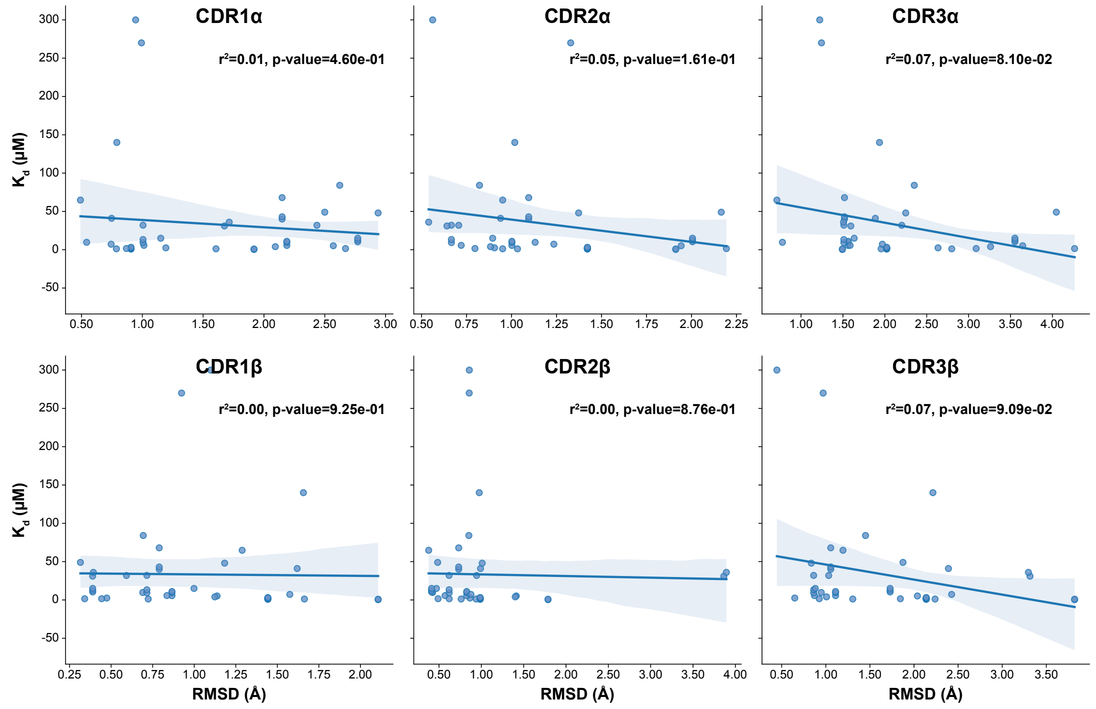


Figure S6: Correlating RMSD changes of CDR loops to affinity.

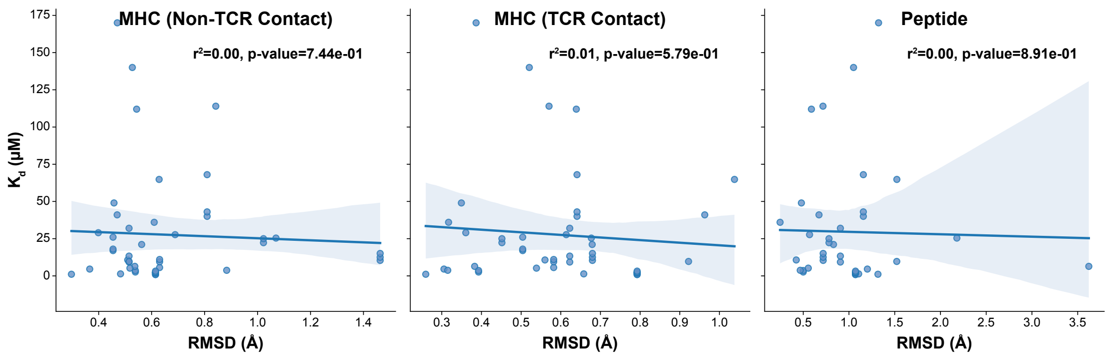


Figure S7: Correlating affinity changes to RMSD.

between conformational change and affinity, the trends are subtle and one is not indicative of the other.

### S2 Correlation of CDR loop length and conformational change

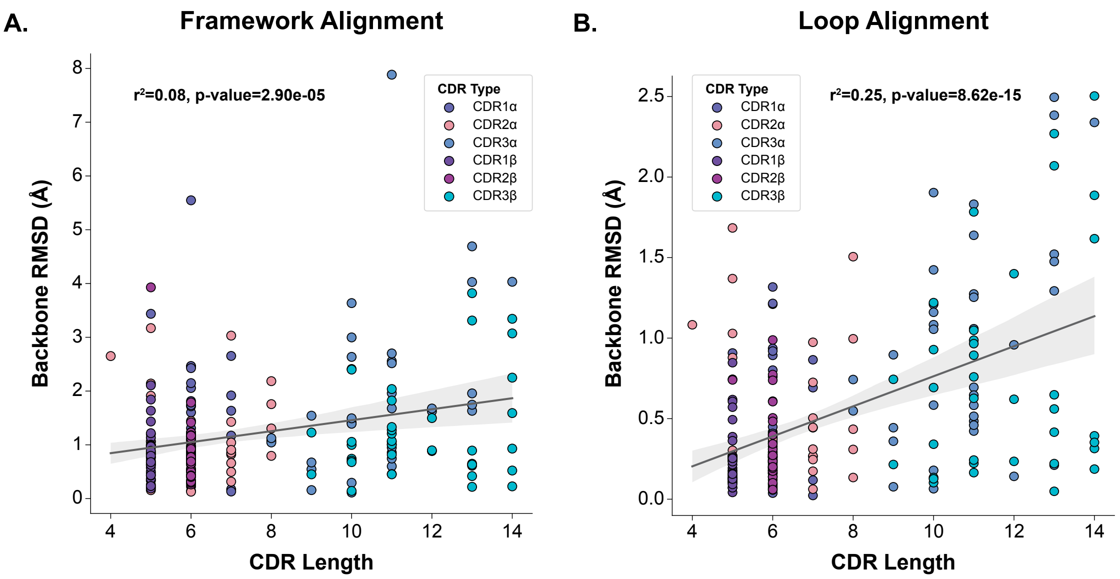


Figure S8: Correlation of CDR loop length to conformational change. **A.** Conformational changes are measured when TCRs are aligned on framework regions. **B.** Conformational chagnes are measured when CDR loops are aligned before comparison.

Probing the underlying causes of conformational change in CDR loops, we investigated the role loop length plays in allowing for conformational change. Shown in Fig. S8A, bulk CDR movements are weakly correlated to RMSD changes between *apo* and *holo* states ($R^2 = 0.08$). However, Fig. S8B shows a stronger correlation to the deformation effect of loops ($R^2 = 0.25$). The increased correlation makes sense since other parts of the protein can drive the conformational changes from the framework regions, but when the loops are aligned together, the only differences can be driven by changes in the loops themselves, implying the loop length has more of an effect. From these results, it seems loop length has some impact on the inherent flexibility of loops, but other factors also likely modulate the flexibility of these loops such as amino acid composition and interactions with other CDR loops [4].

Table S1: Breakdown of the data used in the analysis of *apo* and *holo* forms of TCR and pMHC structures. In total there are 358 structures coming from 255 PDB entries and encompassing 25 unique TCRs, 20 MHC alleles, and 58 peptides.

| PDB ID | Structure Type | α-chain ID | β-chain ID | Antigen Chain ID | MHC Chain ID |
|---|---|---|---|---|---|
| 1bii | pMHC-I | - | - | P | A |
| 1ddh | pMHC-I | - | - | P | A |
| 1duz | pMHC-I | - | - | C | A |
| 1duz | pMHC-I | - | - | F | D |
| 1fzj | pMHC-I | - | - | P | A |
| 1fzm | pMHC-I | - | - | P | A |
| 1hhi | pMHC-I | - | - | C | A |
| 1hhi | pMHC-I | - | - | F | D |
| 1hhk | pMHC-I | - | - | C | A |
| 1hhk | pMHC-I | - | - | F | D |
| 1hoc | pMHC-I | - | - | C | A |
| 1i4f | pMHC-I | - | - | C | A |
| 1jf1 | pMHC-I | - | - | C | A |
| 1kpu | pMHC-I | - | - | P | A |
| 1m05 | pMHC-I | - | - | E | A |
| 1m05 | pMHC-I | - | - | F | C |
| 1s9w | pMHC-I | - | - | C | A |
| 1tvb | pMHC-I | - | - | C | A |
| 1tvb | pMHC-I | - | - | F | D |
| 1tvh | pMHC-I | - | - | C | A |
| 1tvh | pMHC-I | - | - | F | D |
| 1wby | pMHC-I | - | - | C | A |
| 1yn6 | pMHC-I | - | - | C | A |
| 1zhl | pMHC-I | - | - | C | A |
| 2av1 | pMHC-I | - | - | C | A |
| 2av1 | pMHC-I | - | - | F | D |
| 2av7 | pMHC-I | - | - | C | A |
| 2av7 | pMHC-I | - | - | F | D |
| 2bss | pMHC-I | - | - | C | A |
| 2clv | pMHC-I | - | - | C | A |
| 2clv | pMHC-I | - | - | M | H |
| 2clz | pMHC-I | - | - | C | A |
| 2clz | pMHC-I | - | - | M | H |
| 2git | pMHC-I | - | - | C | A |
| 2gt9 | pMHC-I | - | - | C | A |
| 2gt9 | pMHC-I | - | - | F | D |
| 2guo | pMHC-I | - | - | C | A |
| 2guo | pMHC-I | - | - | F | D |
| 2mha | pMHC-I | - | - | E | A |
| 2mha | pMHC-I | - | - | F | C |

Continued on next page

Table S1: Breakdown of the data used in the analysis of *apo* and *holo* forms of TCR and pMHC structures. In total there are 358 structures coming from 255 PDB entries and encompassing 25 unique TCRs, 20 MHC alleles, and 58 peptides.

| PDB ID | Structure Type | α-chain ID | β-chain ID | Antigen Chain ID | MHC Chain ID |
|---|---|---|---|---|---|
| 2vaa | pMHC-I | - | - | P | A |
| 2vll | pMHC-I | - | - | C | A |
| 2vll | pMHC-I | - | - | F | D |
| 2x4r | pMHC-I | - | - | C | A |
| 2x4r | pMHC-I | - | - | F | D |
| 3ecb | pMHC-I | - | - | P | A |
| 3gso | pMHC-I | - | - | P | A |
| 3h7b | pMHC-I | - | - | C | A |
| 3h7b | pMHC-I | - | - | F | D |
| 3h9h | pMHC-I | - | - | C | A |
| 3h9h | pMHC-I | - | - | F | D |
| 3ixa | pMHC-I | - | - | C | A |
| 3ixa | pMHC-I | - | - | F | D |
| 3kpp | pMHC-I | - | - | C | A |
| 3mre | pMHC-I | - | - | P | A |
| 3mrm | pMHC-I | - | - | P | A |
| 3nfn | pMHC-I | - | - | C | A |
| 3pwl | pMHC-I | - | - | C | A |
| 3pwl | pMHC-I | - | - | F | D |
| 3qfd | pMHC-I | - | - | C | A |
| 3qfd | pMHC-I | - | - | F | D |
| 3quk | pMHC-I | - | - | C | A |
| 3quk | pMHC-I | - | - | F | D |
| 3sko | pMHC-I | - | - | C | A |
| 3tbs | pMHC-I | - | - | C | A |
| 3tbs | pMHC-I | - | - | F | D |
| 3tby | pMHC-I | - | - | C | A |
| 3tby | pMHC-I | - | - | F | D |
| 3tby | pMHC-I | - | - | I | G |
| 3tby | pMHC-I | - | - | L | J |
| 3vfm | pMHC-I | - | - | C | A |
| 3vfn | pMHC-I | - | - | C | A |
| 3vfp | pMHC-I | - | - | C | A |
| 3vxn | pMHC-I | - | - | C | A |
| 3vxo | pMHC-I | - | - | C | A |
| 3vxo | pMHC-I | - | - | F | D |
| 3vxp | pMHC-I | - | - | C | A |
| 3vxp | pMHC-I | - | - | F | D |
| 3x13 | pMHC-I | - | - | C | A |
| 4g8i | pMHC-I | - | - | C | A |

Continued on next page

Table S1: Breakdown of the data used in the analysis of *apo* and *holo* forms of TCR and pMHC structures. In total there are 358 structures coming from 255 PDB entries and encompassing 25 unique TCRs, 20 MHC alleles, and 58 peptides.

| PDB ID | Structure Type | α-chain ID | β-chain ID | Antigen Chain ID | MHC Chain ID |
|---|---|---|---|---|---|
| 4g9d | pMHC-I | - | - | C | A |
| 4hux | pMHC-I | - | - | C | A |
| 4jfp | pMHC-I | - | - | C | A |
| 4jfp | pMHC-I | - | - | F | D |
| 4pr5 | pMHC-I | - | - | C | A |
| 4pra | pMHC-I | - | - | C | A |
| 4u1i | pMHC-I | - | - | C | A |
| 4u1j | pMHC-I | - | - | C | A |
| 4wu5 | pMHC-I | - | - | C | A |
| 4wu5 | pMHC-I | - | - | F | D |
| 4wu7 | pMHC-I | - | - | C | A |
| 4wu7 | pMHC-I | - | - | F | D |
| 5hga | pMHC-I | - | - | C | A |
| 5hga | pMHC-I | - | - | F | D |
| 5hgb | pMHC-I | - | - | C | A |
| 5hgb | pMHC-I | - | - | F | D |
| 5hgb | pMHC-I | - | - | I | G |
| 5hgb | pMHC-I | - | - | L | J |
| 5hgd | pMHC-I | - | - | C | A |
| 5hgd | pMHC-I | - | - | F | D |
| 5hgh | pMHC-I | - | - | C | A |
| 5hhp | pMHC-I | - | - | C | A |
| 5xos | pMHC-I | - | - | C | A |
| 6amt | pMHC-I | - | - | C | A |
| 6amt | pMHC-I | - | - | F | D |
| 6at5 | pMHC-I | - | - | C | A |
| 6g9r | pMHC-I | - | - | P | A |
| 6g9r | pMHC-I | - | - | I | C |
| 6g9r | pMHC-I | - | - | J | E |
| 6g9r | pMHC-I | - | - | K | G |
| 6gh1 | pMHC-I | - | - | P | A |
| 6gh1 | pMHC-I | - | - | Q | C |
| 6gh1 | pMHC-I | - | - | R | E |
| 6gh1 | pMHC-I | - | - | Z | G |
| 6jtp | pMHC-I | - | - | C | A |
| 6mt6 | pMHC-I | - | - | B | A |
| 6npr | pMHC-I | - | - | R | A |
| 6npr | pMHC-I | - | - | P | C |
| 6q3k | pMHC-I | - | - | P | A |
| 6ujo | pMHC-I | - | - | C | A |

Continued on next page

Table S1: Breakdown of the data used in the analysis of *apo* and *holo* forms of TCR and pMHC structures. In total there are 358 structures coming from 255 PDB entries and encompassing 25 unique TCRs, 20 MHC alleles, and 58 peptides.

| PDB ID | Structure Type | α-chain ID | β-chain ID | Antigen Chain ID | MHC Chain ID |
|---|---|---|---|---|---|
| 6ujq | pMHC-I | - | - | C | A |
| 6uli | pMHC-I | - | - | C | A |
| 6vr5 | pMHC-I | - | - | P | A |
| 6vr5 | pMHC-I | - | - | Q | D |
| 7l1c | pMHC-I | - | - | C | A |
| 7mkb | pMHC-I | - | - | C | A |
| 7n1a | pMHC-I | - | - | C | A |
| 7n1a | pMHC-I | - | - | F | D |
| 7n1b | pMHC-I | - | - | C | A |
| 7n1b | pMHC-I | - | - | F | D |
| 7n5q | pMHC-I | - | - | C | A |
| 7n5q | pMHC-I | - | - | H | F |
| 7n6d | pMHC-I | - | - | C | A |
| 7n6d | pMHC-I | - | - | G | E |
| 7n6d | pMHC-I | - | - | K | I |
| 7n6d | pMHC-I | - | - | O | M |
| 7n9j | pMHC-I | - | - | C | A |
| 7nmd | pMHC-I | - | - | C | A |
| 7nmd | pMHC-I | - | - | F | D |
| 7ow3 | pMHC-I | - | - | C | A |
| 7ow3 | pMHC-I | - | - | F | D |
| 7ow3 | pMHC-I | - | - | L | J |
| 7ow4 | pMHC-I | - | - | C | A |
| 7ow4 | pMHC-I | - | - | F | D |
| 7ow4 | pMHC-I | - | - | I | G |
| 7p3d | pMHC-I | - | - | C | A |
| 7r7v | pMHC-I | - | - | C | A |
| 7rtd | pMHC-I | - | - | C | A |
| 1kgc | TCR | D | E | - | - |
| 1tcr | TCR | A | B | - | - |
| 2bnu | TCR | A | B | - | - |
| 2pyf | TCR | A | B | - | - |
| 2vlm | TCR | D | E | - | - |
| 3dx9 | TCR | A | B | - | - |
| 3dx9 | TCR | C | D | - | - |
| 3qeu | TCR | A | B | - | - |
| 3qeu | TCR | D | E | - | - |
| 3qh3 | TCR | A | B | - | - |
| 3skn | TCR | A | B | - | - |
| 3skn | TCR | C | D | - | - |

Continued on next page

Table S1: Breakdown of the data used in the analysis of *apo* and *holo* forms of TCR and pMHC structures. In total there are 358 structures coming from 255 PDB entries and encompassing 25 unique TCRs, 20 MHC alleles, and 58 peptides.

| PDB ID | Structure Type | α-chain ID | β-chain ID | Antigen Chain ID | MHC Chain ID |
|---|---|---|---|---|---|
| 3skn | TCR | E | F | - | - |
| 3skn | TCR | G | H | - | - |
| 3tf7 | TCR | i | I | - | - |
| 3tf7 | TCR | k | K | - | - |
| 3utp | TCR | D | E | - | - |
| 3utp | TCR | K | L | - | - |
| 3vxq | TCR | A | B | - | - |
| 3vxq | TCR | D | E | - | - |
| 3vxt | TCR | A | B | - | - |
| 3vxt | TCR | C | D | - | - |
| 4jfh | TCR | D | E | - | - |
| 4qrp | TCR | K | L | - | - |
| 5iw1 | TCR | A | B | - | - |
| 5iw1 | TCR | C | D | - | - |
| 5iw1 | TCR | E | F | - | - |
| 5nmd | TCR | A | B | - | - |
| 5nmd | TCR | C | D | - | - |
| 5yxu | TCR | F | G | - | - |
| 6at6 | TCR | A | B | - | - |
| 6rp9 | TCR | K | L | - | - |
| 6vth | TCR | A | B | - | - |
| 6vth | TCR | D | E | - | - |
| 7amp | TCR | A | B | - | - |
| 7n1c | TCR | D | E | - | - |
| 7n1d | TCR | A | B | - | - |
| 7r7z | TCR | A | B | - | - |
| 8gop | TCR | A | B | - | - |
| 1ao7 | TCR:pMHC-I | D | E | C | A |
| 1bd2 | TCR:pMHC-I | D | E | C | A |
| 1fo0 | TCR:pMHC-I | A | B | P | H |
| 1g6r | TCR:pMHC-I | A | B | P | H |
| 1g6r | TCR:pMHC-I | C | D | Q | I |
| 1mi5 | TCR:pMHC-I | D | E | C | A |
| 1mwa | TCR:pMHC-I | A | B | P | H |
| 1mwa | TCR:pMHC-I | C | D | Q | I |
| 1nam | TCR:pMHC-I | A | B | P | H |
| 1oga | TCR:pMHC-I | D | E | C | A |
| 1qrn | TCR:pMHC-I | D | E | C | A |
| 1qse | TCR:pMHC-I | D | E | C | A |
| 1qsf | TCR:pMHC-I | D | E | C | A |

Continued on next page

Table S1: Breakdown of the data used in the analysis of *apo* and *holo* forms of TCR and pMHC structures. In total there are 358 structures coming from 255 PDB entries and encompassing 25 unique TCRs, 20 MHC alleles, and 58 peptides.

| PDB ID | Structure Type | α-chain ID | β-chain ID | Antigen Chain ID | MHC Chain ID |
|---|---|---|---|---|---|
| 2bnq | TCR:pMHC-I | D | E | C | A |
| 2bnr | TCR:pMHC-I | D | E | C | A |
| 2ckb | TCR:pMHC-I | A | B | P | H |
| 2ckb | TCR:pMHC-I | C | D | Q | I |
| 2f53 | TCR:pMHC-I | D | E | C | A |
| 2f54 | TCR:pMHC-I | D | E | C | A |
| 2f54 | TCR:pMHC-I | K | L | H | F |
| 2gj6 | TCR:pMHC-I | - | - | C | A |
| 2oi9 | TCR:pMHC-I | B | C | Q | A |
| 2ol3 | TCR:pMHC-I | A | B | P | H |
| 2p5e | TCR:pMHC-I | D | E | C | A |
| 2p5w | TCR:pMHC-I | D | E | C | A |
| 2pye | TCR:pMHC-I | D | E | C | A |
| 2vlj | TCR:pMHC-I | D | E | C | A |
| 2vlk | TCR:pMHC-I | D | E | C | A |
| 3d39 | TCR:pMHC-I | D | E | C | A |
| 3d3v | TCR:pMHC-I | D | E | C | A |
| 3dxa | TCR:pMHC-I | D | E | C | A |
| 3dxa | TCR:pMHC-I | I | J | H | F |
| 3dxa | TCR:pMHC-I | N | O | M | K |
| 3gsn | TCR:pMHC-I | A | B | P | H |
| 3h9s | TCR:pMHC-I | D | E | C | A |
| 3hg1 | TCR:pMHC-I | D | E | C | A |
| 3kpr | TCR:pMHC-I | D | E | C | A |
| 3kpr | TCR:pMHC-I | I | J | H | F |
| 3kps | TCR:pMHC-I | D | E | C | A |
| 3o4l | TCR:pMHC-I | D | E | C | A |
| 3pwp | TCR:pMHC-I | D | E | C | A |
| 3qdg | TCR:pMHC-I | D | E | C | A |
| 3qdj | TCR:pMHC-I | D | E | C | A |
| 3qdm | TCR:pMHC-I | D | E | C | A |
| 3qeq | TCR:pMHC-I | D | E | C | A |
| 3qfj | TCR:pMHC-I | D | E | C | A |
| 3sjv | TCR:pMHC-I | D | E | C | A |
| 3sjv | TCR:pMHC-I | I | J | H | F |
| 3sjv | TCR:pMHC-I | N | O | M | K |
| 3sjv | TCR:pMHC-I | S | T | R | P |
| 3tf7 | TCR:pMHC-I | c | C | B | A |
| 3tf7 | TCR:pMHC-I | g | G | F | E |
| 3tfk | TCR:pMHC-I | C | D | B | A |

Continued on next page

Table S1: Breakdown of the data used in the analysis of *apo* and *holo* forms of TCR and pMHC structures. In total there are 358 structures coming from 255 PDB entries and encompassing 25 unique TCRs, 20 MHC alleles, and 58 peptides.

| PDB ID | Structure Type | α-chain ID | β-chain ID | Antigen Chain ID | MHC Chain ID |
|---|---|---|---|---|---|
| 3tjh | TCR:pMHC-I | C | D | B | A |
| 3tpu | TCR:pMHC-I | A | B | J | I |
| 3tpu | TCR:pMHC-I | C | D | F | E |
| 3tpu | TCR:pMHC-I | G | H | L | K |
| 3tpu | TCR:pMHC-I | M | N | R | Q |
| 3uts | TCR:pMHC-I | D | E | C | A |
| 3uts | TCR:pMHC-I | I | J | H | F |
| 3utt | TCR:pMHC-I | D | E | C | A |
| 3utt | TCR:pMHC-I | I | J | H | F |
| 3vxm | TCR:pMHC-I | D | E | C | A |
| 3vxr | TCR:pMHC-I | D | E | C | A |
| 3vxs | TCR:pMHC-I | D | E | C | A |
| 3vxu | TCR:pMHC-I | D | E | C | A |
| 3vxu | TCR:pMHC-I | I | J | H | F |
| 3w0w | TCR:pMHC-I | D | E | C | A |
| 4ftv | TCR:pMHC-I | D | E | C | A |
| 4g8g | TCR:pMHC-I | D | E | C | A |
| 4g9f | TCR:pMHC-I | D | E | C | A |
| 4jfd | TCR:pMHC-I | D | E | C | A |
| 4jfe | TCR:pMHC-I | D | E | C | A |
| 4jff | TCR:pMHC-I | D | E | C | A |
| 4jrx | TCR:pMHC-I | D | E | C | A |
| 4jry | TCR:pMHC-I | D | E | C | A |
| 4l3e | TCR:pMHC-I | D | E | C | A |
| 4ms8 | TCR:pMHC-I | C | D | B | A |
| 4mvb | TCR:pMHC-I | C | D | B | A |
| 4mxq | TCR:pMHC-I | C | D | B | A |
| 4n0c | TCR:pMHC-I | C | D | B | A |
| 4n0c | TCR:pMHC-I | G | H | F | E |
| 4n5e | TCR:pMHC-I | C | D | B | A |
| 4prh | TCR:pMHC-I | D | E | C | A |
| 4prp | TCR:pMHC-I | D | E | C | A |
| 4qok | TCR:pMHC-I | D | E | C | A |
| 4qrp | TCR:pMHC-I | D | E | C | A |
| 4qrp | TCR:pMHC-I | J | I | H | F |
| 5c07 | TCR:pMHC-I | D | E | C | A |
| 5c07 | TCR:pMHC-I | I | J | H | F |
| 5c08 | TCR:pMHC-I | D | E | C | A |
| 5c08 | TCR:pMHC-I | I | J | H | F |
| 5c09 | TCR:pMHC-I | D | E | C | A |

Continued on next page

Table S1: Breakdown of the data used in the analysis of *apo* and *holo* forms of TCR and pMHC structures. In total there are 358 structures coming from 255 PDB entries and encompassing 25 unique TCRs, 20 MHC alleles, and 58 peptides.

| PDB ID | Structure Type | α-chain ID | β-chain ID | Antigen Chain ID | MHC Chain ID |
|---|---|---|---|---|---|
| 5c09 | TCR:pMHC-I | I | J | H | F |
| 5c0a | TCR:pMHC-I | D | E | C | A |
| 5c0a | TCR:pMHC-I | I | J | H | F |
| 5c0b | TCR:pMHC-I | D | E | C | A |
| 5c0b | TCR:pMHC-I | I | J | H | F |
| 5c0c | TCR:pMHC-I | D | E | H | F |
| 5c0c | TCR:pMHC-I | I | J | C | A |
| 5hhm | TCR:pMHC-I | D | E | C | A |
| 5hhm | TCR:pMHC-I | I | J | H | F |
| 5hho | TCR:pMHC-I | D | E | C | A |
| 5hyj | TCR:pMHC-I | D | E | C | A |
| 5hyj | TCR:pMHC-I | I | J | H | F |
| 5isz | TCR:pMHC-I | D | E | C | A |
| 5ivx | TCR:pMHC-I | E | F | P | A |
| 5m00 | TCR:pMHC-I | G | H | P | A |
| 5m01 | TCR:pMHC-I | G | H | P | A |
| 5m02 | TCR:pMHC-I | G | H | P | A |
| 5nht | TCR:pMHC-I | A | B | P | H |
| 5nme | TCR:pMHC-I | D | E | C | A |
| 5nme | TCR:pMHC-I | I | J | H | F |
| 5nmf | TCR:pMHC-I | D | E | C | A |
| 5nmf | TCR:pMHC-I | I | J | H | F |
| 5nmg | TCR:pMHC-I | D | E | C | A |
| 5nmg | TCR:pMHC-I | I | J | H | F |
| 5nqk | TCR:pMHC-I | A | B | P | H |
| 5sws | TCR:pMHC-I | D | E | C | A |
| 5xot | TCR:pMHC-I | D | E | C | A |
| 5yxn | TCR:pMHC-I | A | B | I | C |
| 5yxu | TCR:pMHC-I | A | B | I | C |
| 6am5 | TCR:pMHC-I | D | E | C | A |
| 6amu | TCR:pMHC-I | D | E | C | A |
| 6avf | TCR:pMHC-I | A | B | P | H |
| 6bj2 | TCR:pMHC-I | D | E | C | A |
| 6bj3 | TCR:pMHC-I | D | H | C | A |
| 6d78 | TCR:pMHC-I | D | E | C | A |
| 6dkp | TCR:pMHC-I | D | E | C | A |
| 6eqa | TCR:pMHC-I | D | E | C | A |
| 6eqb | TCR:pMHC-I | D | E | C | A |
| 6g9q | TCR:pMHC-I | G | H | P | A |
| 6mtm | TCR:pMHC-I | D | E | C | A |

Continued on next page

Table S1: Breakdown of the data used in the analysis of *apo* and *holo* forms of TCR and pMHC structures. In total there are 358 structures coming from 255 PDB entries and encompassing 25 unique TCRs, 20 MHC alleles, and 58 peptides.

| PDB ID | Structure Type | α-chain ID | β-chain ID | Antigen Chain ID | MHC Chain ID |
|---|---|---|---|---|---|
| 6q3s | TCR:pMHC-I | D | E | C | A |
| 6rp9 | TCR:pMHC-I | D | E | C | A |
| 6rp9 | TCR:pMHC-I | I | J | H | F |
| 6tmo | TCR:pMHC-I | D | E | C | A |
| 6tro | TCR:pMHC-I | D | E | C | A |
| 6uk2 | TCR:pMHC-I | - | - | C | A |
| 6uk4 | TCR:pMHC-I | - | - | C | A |
| 6uln | TCR:pMHC-I | - | - | C | A |
| 6ulr | TCR:pMHC-I | - | - | C | A |
| 6vm7 | TCR:pMHC-I | - | - | C | A |
| 6vm8 | TCR:pMHC-I | - | - | C | A |
| 6vm9 | TCR:pMHC-I | - | - | C | A |
| 6vma | TCR:pMHC-I | - | - | C | A |
| 6vrm | TCR:pMHC-I | D | E | P | A |
| 6vrn | TCR:pMHC-I | D | E | P | A |
| 6zkw | TCR:pMHC-I | D | E | C | A |
| 7dzm | TCR:pMHC-I | E | D | C | A |
| 7dzn | TCR:pMHC-I | E | D | C | A |
| 7jwi | TCR:pMHC-I | D | E | C | A |
| 7jwj | TCR:pMHC-I | D | E | C | A |
| 7n1e | TCR:pMHC-I | D | E | C | A |
| 7n1f | TCR:pMHC-I | D | E | C | A |
| 7n4k | TCR:pMHC-I | - | - | C | A |
| 7n5c | TCR:pMHC-I | D | E | C | A |
| 7n5p | TCR:pMHC-I | D | E | C | A |
| 7na5 | TCR:pMHC-I | D | E | C | A |
| 7nme | TCR:pMHC-I | D | E | C | A |
| 7nmf | TCR:pMHC-I | D | E | C | A |
| 7ow5 | TCR:pMHC-I | D | E | C | A |
| 7ow6 | TCR:pMHC-I | D | E | C | A |
| 7r80 | TCR:pMHC-I | A | B | E | C |
| 7rrg | TCR:pMHC-I | - | - | C | A |
| 7rtr | TCR:pMHC-I | D | E | C | A |
| 8gom | TCR:pMHC-I | D | E | C | A |
| 8gon | TCR:pMHC-I | D | E | C | A |
| 8gvb | TCR:pMHC-I | A | B | P | H |
| 8gvg | TCR:pMHC-I | A | B | P | H |
| 8gvi | TCR:pMHC-I | A | B | P | H |
